# Quantitative proteomics reveals remodeling of protein repertoire across life phases of *Daphnia pulex*

**DOI:** 10.1101/772442

**Authors:** Leonid Peshkin, Myriam Boukhali, Wilhelm Haas, Marc W. Kirschner, Lev Y. Yampolsky

**Affiliations:** Department of Systems Biology, Harvard Medical School, Boston, MA 02115, USA; Massachusetts General Hospital Cancer Center and Department of Medicine, Harvard Medical School, Building 149, 13th Street, Charlestown, MA 02129; Department of Biological Sciences, East Tennessee State University, Johnson City, TN 31714, USA

**Keywords:** Daphnia, life cycle, Proteomics, trehalose synthesis, vitellogenins

## Abstract

The microcrustacean *Daphnia* is becoming an organism of choice for genomic and proteomic studies of the effects of environmental stressors. However, the changes in protein expression across the life cycle have not been fully characterized. We analyzed the proteomes of adult females, juveniles, asexually produced embryos, and the ephippia - resting stages containing two sexually produced diapausing freezing- and desiccation-resistant embryos. Overall, we were significantly more likely to detect proteins with known molecular functions than proteins with no detectable orthology. Similarly we could detect those with stronger gene model support, as judged by mutual best BLAST hits between two independent genome assemblies than those without such support. This suggests that we could apply our proteomics pipeline to verify hypothesized proteins, even given less-than-perfect reference gene models. In particular, we observed up-regulation of vitellogenins and down-regulation of actins and myosins in embryos of both types, as compared to juveniles and adults and overrepresentation of cell-cycle related proteins in the developing embryos, as compared to both diapausing embryos and adults. We found upregulation of small heat-shock proteins and redox peroxidases, as well as overrepresentation of stress-response proteins in the ephippium relative to the asexually produced non-diapausing embryos. The ephippium also showed up-regulation of three trehalose-synthesis proteins and down-regulation of a trehalose hydrolase, consistent with the role of trehalose in protection against freezing and desiccation.

**Statement of significance of the study:** Freshwater plankton crustacean *Daphnia* is rapidly becoming a model organism of choice for ecological and developmental genomics. While there have been several advances towards establishing the protocols and reference datasets for proteomics, a detailed dataset covering several main steps of asexual and sexual phases of *Daphnia* life cycle is not yet available. Moreover, different versions of *D. pulex* genome differ in the number of protein-coding genes identified; it is unclear whether these differences are caused by differences between sequenced genotypes or between gene model methodology used. In this study we report LC-MS2/MS3 proteomes of whole body adult females, juvenile females, asexually produces embryos and diapausing eggs capable of surviving freezing and desiccation.

## Introduction

MS-based proteomics is a major tool for understanding changes in gene expression, biochemistry, and physiology both during development and the response to environmental change. Microcrustacean *Daphnia*, an emerging model organism for biomedical research, has several useful features allowing characterization of life-history and physiological plasticity, including clonal reproduction, short lifespan and well-characterized genome. Here we report proteome differences between parthenogenetically produced embryos, juveniles, adults and sexually produced resting eggs that can be used as a base-line reference for protein expression analysis across the life cycle of an organism. Previous advances in *Daphnia* proteomics identified difficulties in obtaining a large number of intact and identifiable proteins due to high protease activity in tissue extracts[1–3], often resulting in a relatively small number of proteins reported (often 500-1500 out of over 25,000 predicted protein coding genes; the highest number reported close to 4000 [4].

Despite these difficulties, a variety of proteomic responses have been reported, including responses to various ecological stimuli, such as temperature [5,6], food availability[6], hypoxia[7], microgravity[8], and presence of predators[9–11] or toxicants[4, 12–14]. Other studies focused on characterization of specific groups of proteins, such as neuropeptides and hormones[15]. Yet, *Daphnia* proteome remains largely uncharacterized[3], including the lack of proteomics data across different life cycle phases.

In favorable conditions *Daphnia* reproduce asexually throughout the life cycle producing subitaneous eggs developing into offspring genetically identical to each other and the mother[16]. These eggs undergo immediate development in the female’s brood chamber and are released as neonates largely resembling adults. In response to environmental cues such as population density, temperature, and photoperiod meiosis is induced and haploid resting eggs are produced, each sealed in protective structure called *ephippium*. These eggs require fertilization to survive; they arrest development in early gastrulation phase and are reactivated after a period of exposure to darkness and low temperatures[16].

Several *a priori* predictions can be made about proteins expressed during each phase of the life cycle. Embryos, both asexually and sexually produced, are expected to contain higher levels of vitellogenins (glycolipoproteins that are precursors of yolk lipo- and phosphoproteins in many animals including *Daphnia*[17]). Conversely, ephippia embryos that have less or no muscle tissue would be expected to contain less actin and myosin. Diapausing eggs capable of surviving freezing, heating and desiccation are hypothesized to contain high levels of trehalose, a disaccharide commonly used in crustaceans and insects as freezing and desiccation protectant [18,19], thus one might expect upregulation of trehalose synthesis and downregulation of trehalose breakdown proteins in ephippia. One may also expect elevated expression in ephippia of stress-tolerance related proteins, such as small heat-shock proteins [20,21] and proteins involved in antioxidant pathways, such as superoxide dysmutases (SODs), peroxidases and other glutathione metabolism enzymes, and thioredoxins[22]. Additionally, desiccation resistant mosquito shows up-regulation of protein repair enzymes protein-L-isoaspartate-(D-aspartate) O-methyltransferases (PIMTs) during desiccation stress [22]. A *Daphnia magna* ortholog of mosquito PIMT has been shown to be up-regulated in freshly laid ephippia (Y.Galimov and O.Gusev, personal communication). On the other hand, parthenogenetically produced embryos are not capable of surviving freezing or desiccation, so these expectations do not apply to them.

Although *Daphnia pulex* genome is relatively well characterized[23, 24], the exact number of protein coding genes in the genome is still subject of debate, as it varies significantly between closely related genomes and/or between gene model methodologies[24]. Embarrassingly, only about 1/3 of protein coding genes in either genome assembly are each other’s mutual best BLAST hits (M.Pfrender, personal communication; L. Yampolsky, unpublished). We used our proteomics data to evaluate the agreement (or lack thereof) between the two genome assemblies using the set of proteins observed in this study as a positive control set likely to be enriched in true positives, i.e., genes that are actually expressed.

## Materials and methods

### Sample Preparation

Adult and juvenile *Daphnia* as well as subitaneous (asexually produced non-diapausing) embryos and resting eggs (ephippia) were sampled from a mass culture of *D.pulex* “TCO” clone. In order to roughly match the amount of protein we used 3 adult animals, 10 juvenile and 30 embryos as well as 11 ephippia per sample, resulting in 3 adult females (over 15 days old), 3 juvenile (3-5 days old), and 3 embryonic samples (within 2 days after egg-laying), and a single ephippium sample. Adults and juveniles were sampled from a mass culture; the subitaneous embryos were dissected from the same females used for the "adult" samples. The ephippia were collected from the bottom of the culture tank within 7 days after fertilization/shedding by the female. The ephippia were homogenized whole, thus containing both embryos and maternal tissues. The single ephippia sample available makes all the comparisons that involve this un-replicated sample preliminary in nature as they are severely affected by likely variation among biological specimens and prep procedures. However, at least for the overrepresentation of GO terms analysis (see below), this was partly ameliorated by downstream statistical analysis eliminating any GO terms associated wit false-positives caused by lack of replication with the ephippial embryos category.

Following Refs[25,26], *Daphnia* were lysed in a buffer containing 75 mM NaCl, 3 % SDS, 1 mM NaF, 1 mM beta-glycerophosphate, 1 mM sodium orthovanadate, 10 mM sodium pyrophosphate, 1 mM PMSF and 1X Roche Complete Mini EDTA free protease inhibitors in 50 mM HEPES, pH 8.5. Lyses was supported by using a pellet pestle. Lysates were then sonicated for 5 minutes in a sonicating water bath before cellular debris was pelleted by centrifugation at 14000 rpm for 5 minutes. Proteins were then reduced with DTT and alkylated with iodoacetamide as previously described and precipitated via methanol-chloroform precipitation [25]. Precipitated proteins were reconstituted in 300 μL of 1 M urea in 50 mM HEPES, pH 8.5 and digested in a two-step process starting with an overnight digest at room temperature with Lys-C (Wako) followed by six hours of digestion with trypsin (sequencing grade, Promega) at 37 ° C. The digest was acidified with TFA and peptides were desalted with C_18_ solid-phase extraction (SPE) (Sep-Pak, Waters) as previously described[25]. The concentration of the desalted peptide solutions was measured with a BCA assay, and peptides were aliquoted into 50 μg portions, which were dried under vacuum and stored at −80 °C until they were labeled with 10-plex TMT reagents (Thermo Scientific) as described previously[25]. Pooled samples were desalted via C_18_ SPE on Sep-Pak cartridges as described above and subjected to basic pH reversed-phase liquid chromatography (bRPLC)[25] over a 4.6 mm x 250 mm ZORBAX Extend C_18_ column (5 μm, 80 Å, Agilent Technologies) with concatenated fraction combining as previously described, and 12 fractions were subjected to quantitative proteomics analysis.

### Liquid Chromatography Mass Spectrometry

All LC-MS2/MS3 experiments were conducted on an Orbitrap Fusion (Thermo Fisher Scientific) coupled to an Easy-nLC 1000 (Thermo Fisher Scientific) with chilled autosampler. Peptides were separated on an in-house pulled, in-house packed microcapillary column (inner diameter, 100 μm; outer diameter, 360 μm). Columns were packed to a final length of 30 cm with GP-C_18_ (1.8 μm, 120 Å, Sepax Technologies). Peptides were eluted with a linear gradient from 11 to 30 % ACN in 0.125 % formic acid over 165 minutes at a flow rate of 300 nL/minute while the column was heated to 60 °C. Electrospray ionization was achieved by applying 1500 V through a stainless steel T-junction at the inlet of the microcapillary column. The Orbitrap Fusion was operated in data-dependent mode, with a survey scan performed over an m/z range of 500-1,200 in the Orbitrap with a resolution of 6×10^4^, automatic gain control (AGC) of 5 × 10^5^, and a maximum injection time of 100 ms. The most abundant ions detected in the survey scan were subjected to MS2 and MS3 experiments to be acquired in a 5 seconds experimental cycle. For MS2 analysis, doubly charged ions were selected from an m/z range of 600-1200, and triply and quadruply charged ions from an m/z range of 500-1200. The ion intensity threshold was set to 5×10^4^ and the isolation window to 0.5 m/z. Peptides were isolated using the quadrupole and fragmented using CID at 30 % normalized collision energy at the rapid scan rate using an AGC target of 1 × 10^4^ and a maximum ion injection time of 35 ms. MS3 analysis was performed using synchronous precursor selection (SPS)[27,28]. Up to 6 MS2 precursors were simultaneously isolated and fragmented for MS3 analysis with an isolation window of 2.5 m/z and HCD fragmentation at 55 % normalized collision energy. MS3 spectra were acquired at a resolution of 5×10^4^ with an AGC target of 5 × 10^4^ and a maximum ion injection time of 86 ms. The lowest m/z for the MS3 scans was set to 110.

### Data Processing and Analysis

Data were processed using an in-house developed software suite [29]. MS2 data were annotated using the Sequest algorithm[30] to search the to the set of *Daphnia pulex* proteins present in UniProt database [31] with the addition of known contaminants such as trypsin, and a target-decoy database strategy was applied to measure false-discovery rates of peptide and protein identifications (Supplementary File 1). Searches were performed with a 50 ppm precursor mass tolerance; 10-plex TMT tags on lysine residues and peptide n-termini (+229.162932 Da) and carbamidomethylation of cysteines (+57.02146 Da) were set as static modifications and oxidation of methionine (+15.99492 Da) as a variable modification. Data were filtered to a peptide and protein false discovery rate of less than 1 % using the target-decoy search strategy[32]. MS2 assignments were filtered using linear discriminant analysis using a combined score generated from the following peptide and spectral properties: XCorr, dCn, number of missed tryptic cleavages, mass deviation, and peptide length (see Supplementary File 2)[29]. The probability of an incorrect peptide annotation was calculated using a posterior error histogram from sorting peptide annotations from the forward and the reversed database based on their LDA score[29]. The probability of an incorrect protein assignment was calculated by multiplying the LDA probabilities of all peptides assigned to the protein[29]. Peptides that matched to more than one protein were assigned to that protein containing the largest number of matched redundant peptide sequences following the law of parsimony [29]. TMT reporter ion intensities were extracted from the MS3 spectra selecting the most intense ion within a 0.003 m/z window centered at the predicted m/z value for each reporter ion and spectra were used for quantification if the sum of the S/N values of all reporter ions was ≥ 100 and the isolation specificity for the precursor ion was ≥ 0.5. Protein intensities were calculated by summing the TMT reporter ions for all peptides assigned to a protein. Intensities were first normalized by the average intensity across all samples relative to the median average across all proteins. In a second normalization step protein intensities measured for each sample were normalized by the average of the median protein intensities measured across the samples. Principal Components analysis was conducted in R using PCA function of FactoMineR 1.40 package[33] with data scaled to unit variance. Biological Processes (BP) GO terms overrepresentation analysis was conducted by PANTHER14.1 gene ontology tool (http://www.pantherdb.org; Ref[34]) using *D. pulex* UniProt annotations and lists of IDs that included proteins showing differences among life stages with FDR<0.01; FDR for Fisher’s Exact Test for overrepresentation set to 0.05.

## Results

### Outcomes of MS analysis and protein identification

Proteomic analysis revealed 34158 peptides that mapped to unique 5099 proteins in the UNIPROT reference. Of these, 5062 proteins were matched to *D.pulex* TCO annotated proteins (version jgi060905 [22]. The likelihood of discovering a protein using our protocol was strikingly non-uniform across molecular function (MF) categories (Table 1). Specifically, proteins detected were highly enriched among enzymes, structural proteins, transporters and RNA-binding proteins; in these categories we detected between one third and one half of all annotated proteins. In contrast, we detected fewer than the expected number of transcription factors and, notably, fewer of the proteins with unknown functions, which were identified by the genome gene model analyses. Here were only able to identify 4%. These proteins were also represented by the fewest peptides (2.89 ±1.16), significantly less than the yield of transporters 13.73 ± 4.24), structural proteins (14.38 ±2.23) and metabolic enzymes (6.64 ±0.24); ANOVA on log-transformed peptide numbers per protein: d.f.=8,5089; F=34.4; P<2.5E-53; Tukey test for the difference between unknown proteins and transporters, structural proteins, and metabolic enzymes: P<0.001).

**Table 1.** Enrichment of molecular function categories among proteins discovered by MS. (** - P<0.01; **** - P<0.0001; individual cells Chi^2^ tests).

| Molecular function | Obs | Exp | P | portion<br>discovered | Number of<br>peptides (SE) |  |
| --- | --- | --- | --- | --- | --- | --- |
| Enzyme | 2598 | 1443.1 | **** | 0.29 | 6.64 | (0.24) |
| Structural | 243 | 87.7 | **** | 0.44 | 14.38 | (2.23) |
| Transporter/channel | 272 | 185.6 | **** | 0.24 | 13.73 | (4.24) |
| Receptor | 100 | 102.0 | n.s. | 0.16 | 4.02 | (0.62) |
| TF | 30 | 70.6 | **** | 0.07 | 2.63 | (0.65) |
| Other DNA binding | 43 | 29.7 | ** | 0.23 | 3.88 | (0.71) |
| RNA binding | 43 | 13.2 | **** | 0.52 | 4.53 | (0.62) |
| Other | 1189 | 825.2 | **** | 0.23 | 6.64 | (0.24) |
| Unknown | 580 | 2340.9 | **** | 0.04 | 2.89 | (1.16) |
| Total | 5098 |  |  | 0.16 | 6.70 | (0.38) |

We also observed that proteins identified as mutual best BLAST hits between the TCO protein reference used in this analysis and a more recent PA42 *D.pulex* protein reference[24] are highly enriched among the proteins we were able to detect. Out of 11697 such proteins we detected 4312 (expected=2004), while among the 18063 proteins with ambiguous hits or no hits we detected only 786 (expected=3094; Fisher Exact Test P<1E-248). This enrichment remains highly significant when analyzed within each MF category.

### A priori expected enrichment of molecular functions across life stages

Normalized signals mapped to individual proteins showed good concordance among biological replicates and these served to convincingly separate the samples on the plane of the first two principal components (Fig.1) with PC1 largely separating juveniles and adults from embryos and PC2 – the ephippium sample from asexual individuals of any life cycle phase. Functional genomics analysis revealed that proteins down-regulated in embryos relative to adults were significantly enriched in enzymes and underrepresented among transcription factors and other DNA-binding proteins, RNA-binding proteins and proteins with “other” molecular functions (Table 2). In addition, in the ephippium sample, in the comparison to subitaneous embryos, proteins with unknown molecular function were overrepresented in the subset of up-regulated proteins. Other molecular functions showed no significant enrichment among proteins differentially expressed in samples representing different stages of the life cycle.

**Fig. 1.**
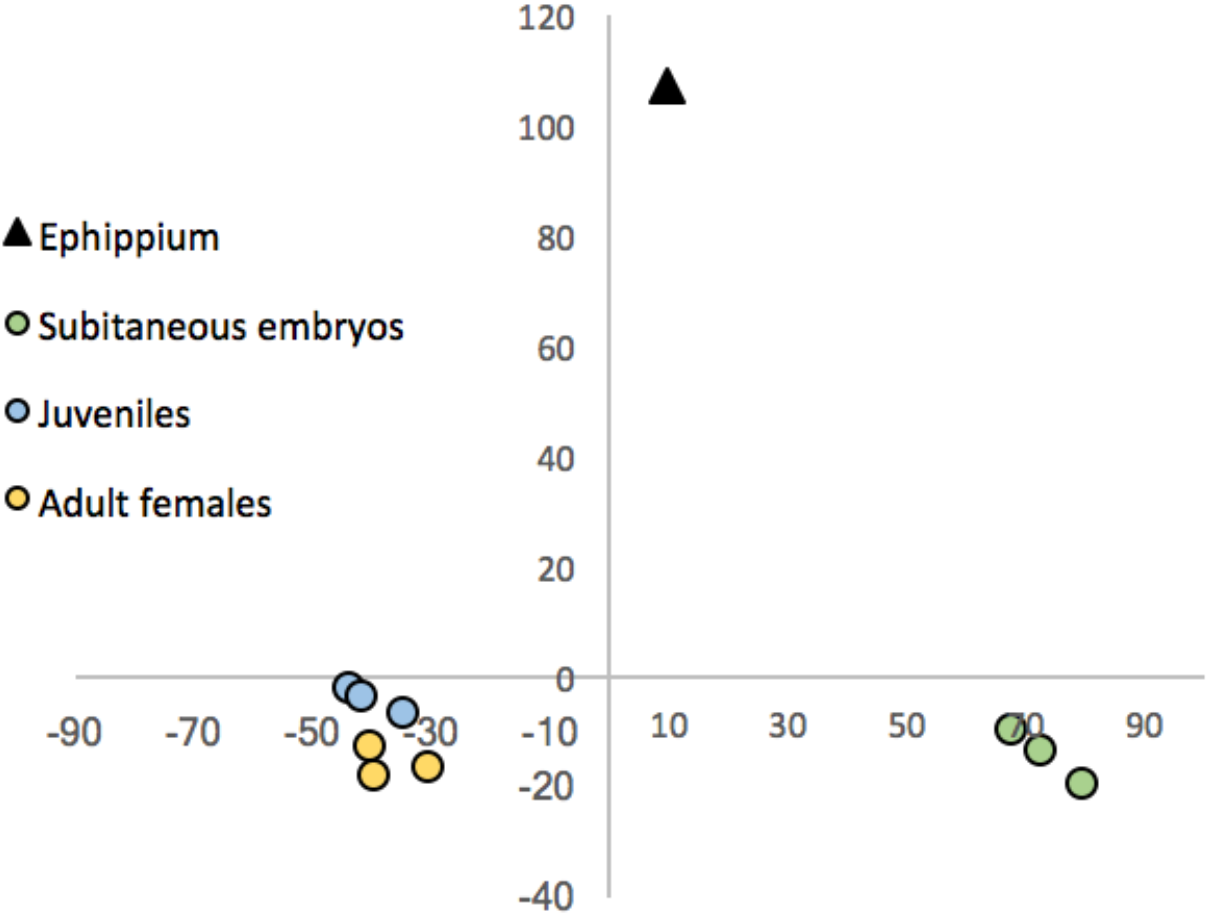
Ten samples analyzed on the plane of the first two principal components explaining 47.1 and 24.4% of variance in protein-specific signals.

**Table 2.**
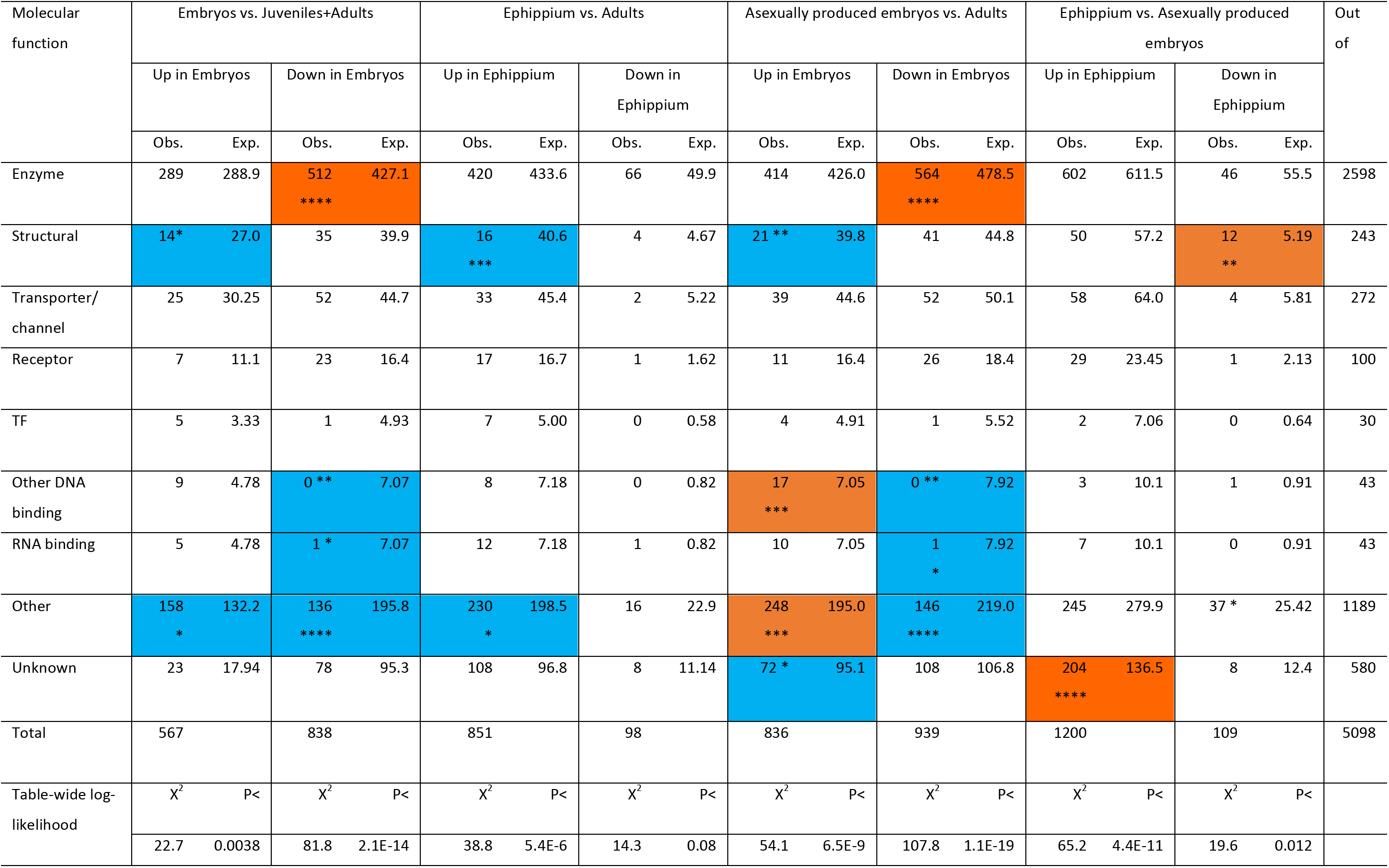
Enrichment analysis of proteins of 7 common molecular functions (plus the “other” and “unknown” categories) among proteins up- or down-regulated (FDR<0.01) in comparisons between embryos and adults as well as between the ephippium and asexually produced embryos. Orange: enriched categories, blue: under-represented categories. P-values (cell’s X^2^): * - <0.025; ** - <0.01; *** - <0.001; **** - <0.0001 (*** and ****: survives table-wide Bonferroni adjustment with P<0.05 and P<0.01, respectively). Bottom: table-wide log-likelihood contingency analysis.

Majority of proteins showing significant differences between adults and embryos tended to be highly expressed in adults, but show low abundance in the embryos (Fig. 2A, red dots). However, there were also a number of proteins up-regulated in the embryos. Several functional groups of proteins *a priori* expected to show differences among life cycle classes showed, as a group, the expected behavior (Fig. 2 B): small HSPs, antioxidant pathways proteins (except thioredoxins), and vitellogenins were more abundant in the embryos than in adults with juveniles showing intermediate values. Unexpectedly, actins and myosins were equally abundant in the embryos juveniles and adults on average (Fig. 2 B), although individual actins and myosins were slightly enriched among genes with a significant down-regulation in embryos (Fisher’s Exact Test two-tailed P<0.035). A similar enrichment among proteins up-regulated in embryos was observed for the antioxidant pathway proteins (Fisher’s Exact Test two-tailed P<0.031).

**Fig. 2.**
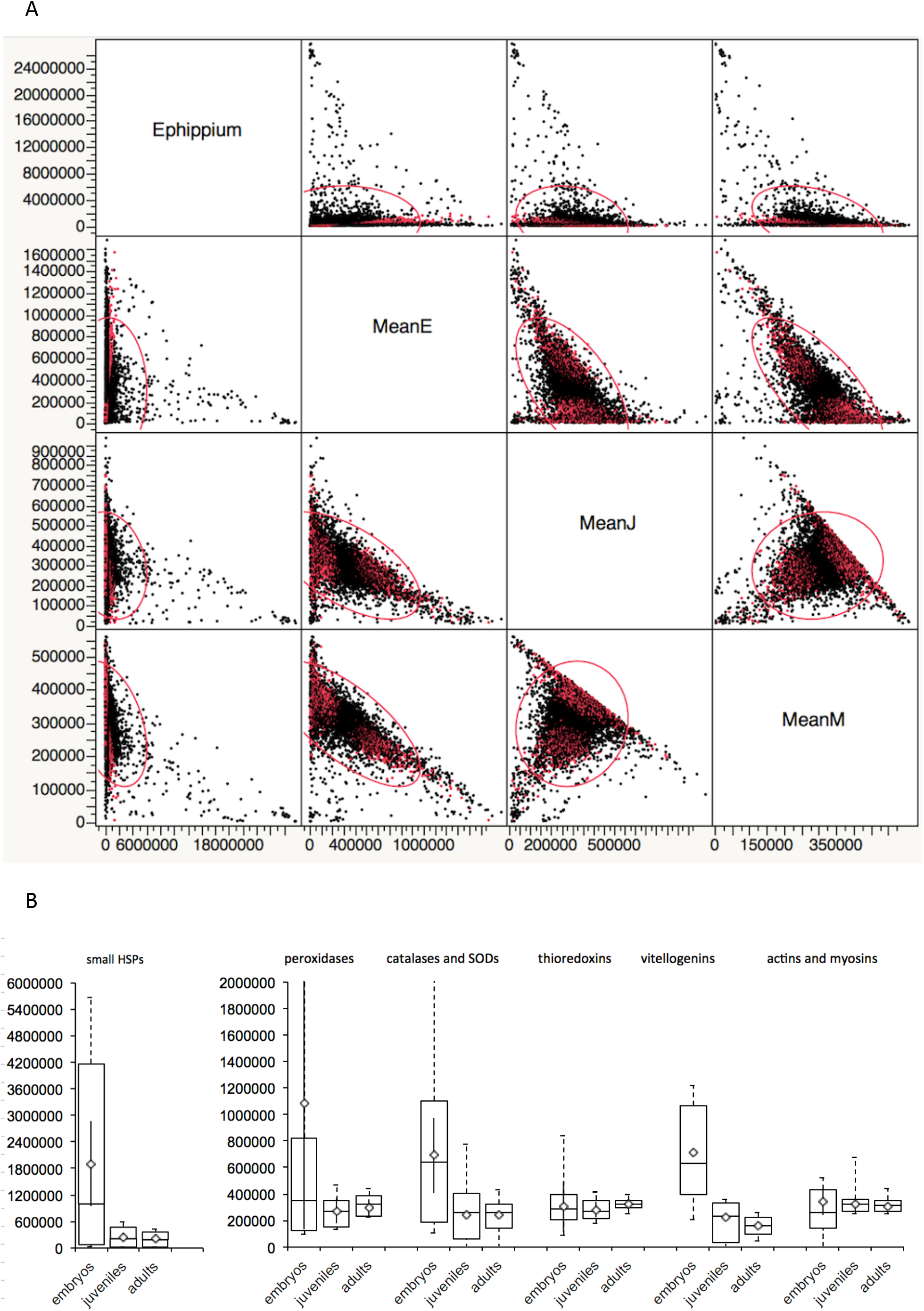
A: scatterplot of relative abundances of 5098 detected proteins in the ephippium (Eph), asexually produced embryos (MeanE), juveniles (MeanJ) and adult females (MeanM). The values are median abundances averaged across replicates. Red: proteins with FDR of the t-test of all empbyos combined vs. adult females <0.01. B: Relative abundances of several fucntional groups of proteins a priori expected to differ among life cycle stages. Boxplots represent mininums, quartiles and maximum across all proteins within the functional category, diamons and solid vertical lines – means and standard errors across proteins. Note a different scale for small HSPs.

The enrichment analysis in comparison between the ephippial sample and subitaneous embryos also revealed a significantly higher occurrence of HSPs and chaperons and antioxidant pathways proteins among proteins up-regulated in the ephippial sample (Fisher’s Exact Test two-tailed P<0.001 and P<0.0002, respectively).

Small HSPs were enriched among proteins up-regulated in the ephippium (Table 2, Fig. 3C) with some showing 10-50 times lower abundance in the subitaneous embryos (Fig. 3C; although one small HPS showed an inverse pattern). Although not significantly enriched (due to the low number of protein in the pathway), the trehalose metabolism genes all show differential expression in the ephippium, three alpha-alspha-trehalose-6-phosphate synthases (that catalyze the rate-limiting step in trehalose synthesis pathway[35], are up-regulated in the ephippium (Fig.2 D; FDR<0.01), while the trehalose hydrolase is down-regulated (FDR<0.01). Similarly, there were numerous peroxidases and small heat shock proteins with higher relative abundance in the ephippium than in the non-diapausing embryos (Fig.2 B,C). Finally, the only PIMT ortholog detected in our samples (UniProt: E9G8Q2) had approximately 3-5-fold higher abundance than either in subitaneous embryos or maternal tissues (FDR = 0.022 and FDR = 0.055, respectively).

**Fig. 3.**
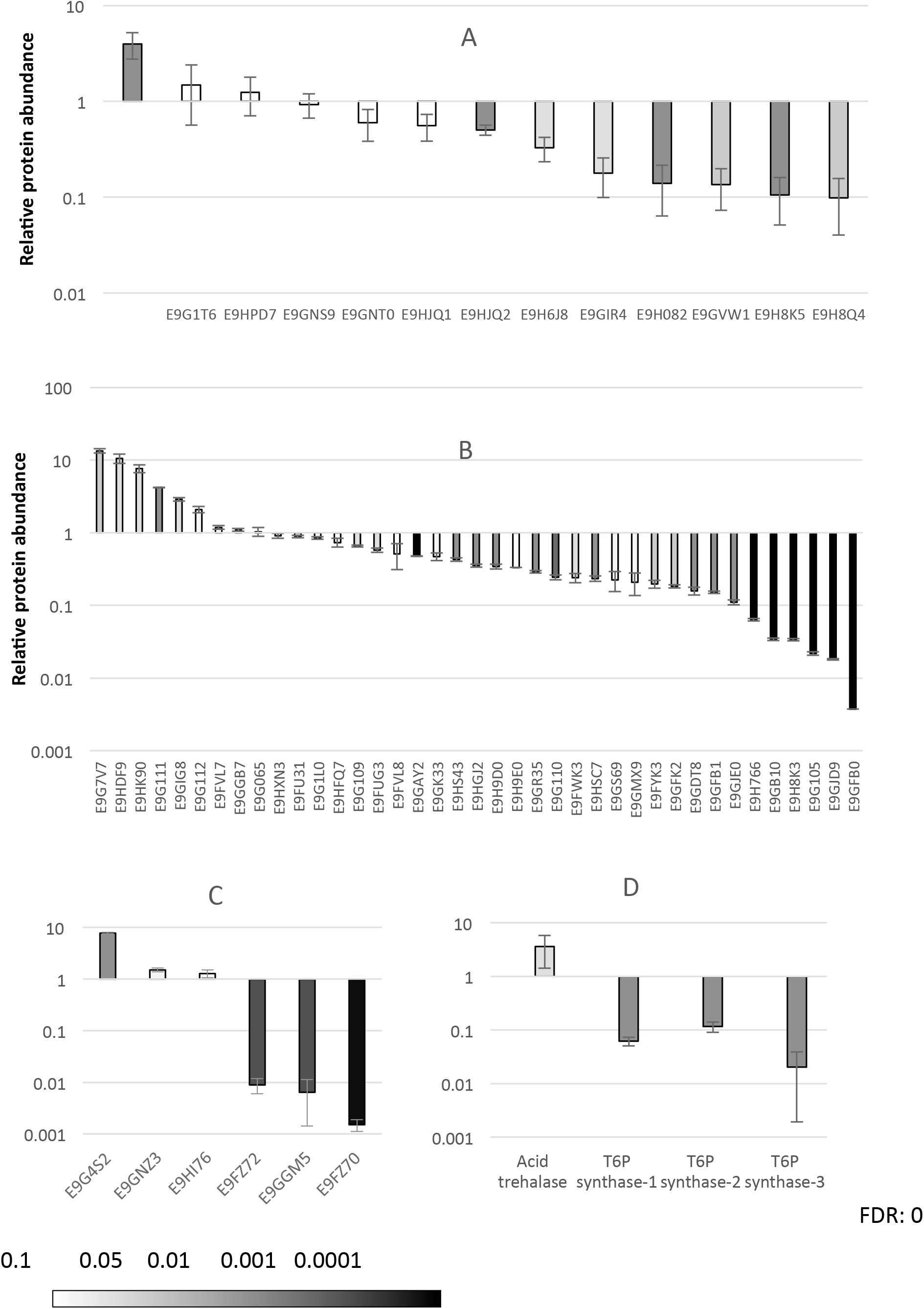
Relative abundance of select proteins in adults vs. embryos and/or the ephippium. A – vitellogenins in juveniles and adults relative to embryos of both types; B – peroxidases in the subitaneous embryos relative to ephippium; C – small HSPs in the subitaneous embryos relative to ephippium; D – trehalose metabolism proteins in the subitaneous embryos relative to ephippium. A: average in all embryos is set to 1; B-D: ephippium is set to 1. White to black shades indicate individual proteins FDR for differential expression test.

### Biological processes overrepresentation across life stages

Proteins with higher abundance in subitaneous (developing) embryos relative to adults showed overrepresentation of a variety of biological processes related to cell division (Supplementary Table 1, Supplementary Fig. 1). These included DNA replication (11-fold enrichment, FDR<10-7) microtubule nucleation and polymerization (both 24-fold overrepresentation and FDR<10-3), mitotic phase transition (12-fold overrepresentation, FDR<10-3). Likewise overrepresented were transcription and ribosome-building PBs indicating active gene expression, as well as histone acetylation, possibly related to ongoing differentiation.

In contrast, among the proteins with higher abundance in adults relative to embryos, the overrepresented biological processes included largely catabolic, respiratory and oxidation-reduction processes (Supplementary Table 2), such as tricarbonic acid cycle (20-fold overrepresentation, FDR<10-8), proton transmembrane transport (12-fold overrepresentation, FDR<10-8), oxidative phosphorylation (9-fold overrepresentation, FDR<10-6), pyruvate metabolism (22-fold overrepresentation, FDR<10-6), Cell adhesion was another BP category overrepresented (5-fold overrepresentation, FDR<10-6). Among the BPs underrepresented in this category of proteins there is, notably, DNA repair pathways (expected 6.54, observed 0, FDR<0.05).

Similar results were obtained in the comparison of both types of embryos to both adults and juveniles combined (Supplementary Tables 3 and 4). The only exception was cell-cycle related proteins that were not overrepresented in this comparison (because there was no active cell division in the arrested embryos in the ephippia, where there was cell division in juveniles).

The overrepresentation analysis of BPs among the proteins with elevated abundance in the single ephippium sample relative to subitaneous embryos revealed, surprisingly, an elevated metabolic activity in the resting embryos than in actively developing ones (Supplementary Table 5, Supplementary Fig. 2). Overrepresented BPs included metabolism of nucleotides, carbohydrates, lipids, pyruvate, amino sugars and aminoglycans (3- to 16-fold overrepresentation). More expectedly, overrepresented were proteins involved in response to stress, including response to reactive oxygen species (7-fold overrepresentation, FDR<0.05), and toxicants (8-fold overrepresentation, FDR<10-8). Likewise, cell adhesion, extracellular matrix organization and intracellular signal transition BPs were overrepresented in the ephippium sample (2- to 5-fold overrepresentation).

In contrast, in the subset of proteins with low abundance in the ephippium sample relative to the subitaneous embryos we observed overrepresentation of nucleotide-, ribonucleoside monophosphate, and DNA biosynthesis processes (11- to 51-fold overrepresentation), translation and protein folding (6- to 33-fold overrepresentation), as one would expect for embryos with arrested development and growth (Supplementary Table 6).

Finally, in the comparison of adult females to juveniles no individual protein achieved FDR<0.01; relaxation of this criterion to FDR<0.05 results in 17 proteins with higher relative abundance in the adults, in which no BPs were significantly overrepresented and 58 proteins with higher relative abundance in juveniles, among which proteins involved in sister chromatid segregation showed a 37-fold overrepresentation (FDR<0.05).

Notably the “unclassified” category is underrepresented in all these comparisons. It would be difficult to hypothesize that constitutively expressed proteins are, on average, less well characterized or less conserved across animals. Rather, we suspect that the Unclassified category is more likely to include false positives or low expression / high noise proteins.

## Discussion

### Proteomics

Despite previous reports that quantitative proteomics in daphnia presents significant challenges [2,3], we were able to obtain a deep coverage of the proteins expressed across very different states of life cycle and conditions, using our standard proteomic sample preparation protocols and analysis procedures and identifying over 5000 proteins. The only previous study that was nearly as successful in identifying individual *D.pulex* proteins [4] utilized the label-free MS protocol. There is a well established tradeoff between the depth of coverage and accuracy of quantitation when comparing label-free and labeled mass spectrometry proteomics analysis with that label free protocol allowing deeper coverage given the same running time per sample, but has a significantly higher variance among samples [36]. Thus, this study reports so far the most successful *Daphnia* proteomics results in terms of total number of proteins identified and in terms of balance between depth and reliability of quantification. The quality of the analysis in proteomics critically depends on the availability of a good reference set of proteins, which has been a challenge for *Daphnia* so far. Our raw data can be easily re-analyzed once a better reference set is available from future DNA sequence analysis.

### Inference for genomic reference

Success of protein identification by MS critically depends on the quality of protein reference used. The reference genome of the clone used in this experiment (TCO) includes a significantly larger number of gene models[23] than a more recently published reference from a closely related member of *D.pulex* group[24]. Our results point to a significant contamination of these gene models with false positives. We were much more likely to detect proteins with known molecular functions (enzymes, structural proteins, transporters and RNA-binding proteins in particular) than those whose molecular function is unknown (Table 1). Of course this can be partially explained by the fact that proteins with low expression are more likely not to be assigned any known molecular function; additionally such proteins also include Daphnia-specific ones with no detectable homology in better annotated genomes. The lower number of peptides by which proteins with unknown MF have been identified may be indicative of their lower expression. Yet, it is also likely that we failed to detect at least some of these proteins because they are artifacts of false gene models. Indeed, the highly significant enrichment of mutual best BLAST hit proteins between the two independently sequenced, assembled and annotated genomes (among proteins that we were able to detect) suggests that low detection success is enriched in proteins with less reliable independent protein or RNA confirmation.

### Functional differences between life cycle stages

We observed several *a priori* expected instances of differential expression between embryos and adults and between sexually and asexually produced embryos, which are, respectively, dormant and subitaneous. These signal include upregulation of vitellogenins and down-regulation of the muscle proteins, actin and myosins in the embryos and up-regulation of several stress resistance-related groups of proteins in the diapausing embryos, including small HSPs, peroxidases, trehalose synthesis enzymes and the single PIMT ortholog, consistent with the previous studies on desiccation-resistant organisms [22]. While overall expression of all peroxidases combined slightly increased along the life cycle (Fig. 2B), there was evidence of numerous individual peroxidases that showed ephippium-specific expression (Fig. 3B). Similarly, the overrepresentation analysis of BPs revealed overrepresentation of proteins involved in response to stress among those with higher relative abundance in the single ephippium sample. Furthermore, we observed a significant overrepresentation of proteins implicated in DNA replication- and mitosis-related BPs in subitaneous embryos relative to both the adults and the single ephippia sample (Supplementary Tables 1, 5, 6; Supplementary Fig. S1). This is consistent with the hypothesis that adults *Daphnia* are largely postmitotic (despite continued growth and molting throughout the lifespan) and any DNA replication may be limited to tissue undergoing endopolyploidization [16]. Overrepresentation of catabolism and respiration processes among proteins with higher abundance in adults (Supplementary Table 2) is consistent with higher energetic expenditures in adults relative to embryos.

Although the lack of replication within the diapausing embryos prevented us for more in-depth analysis, it is clear that the procedure we employed is capable of detecting life cycle differences. The paucity of DNA-binding proteins among those up-regulated in embryos, puzzling at the first glance, can be readily explained by lower number of nuclei in the embryonic samples than in a comparable amount of adult tissues, leading to undersampling of proteins with nuclear localization. Overall, uncharacterized proteins were significantly underrepresented in any BP gene ontology comparison. However, a significant over-representation of proteins with unknown MFs among those that are upregulated in the diapausing embryos was observed. It indicates that those proteins with no assignable ontology that can be identified by proteomics are likely to include diapause-specific proteins, including, possibly, yet unknown proteins, functionality related to adaptations for freezing-, heat- and desiccation tolerance.

## Supporting information

SupplementaryFile1-Proteins

SupplementaryFile3-Peptides

## Data availability

Proteomics data deposited into MassIVE Repository (https://massive.ucsd.edu/), accession number MSV000083711. Relative abundance data by protein and by peptides are available in Supplementary File 1 and 2, respectively.

## Author Contributions

Conceptualization: LP, LY, MWK; Methodology: LP, WH, MWK; Experiments: LP, MB Formal Analysis: LP, WH,LY, Resources: LP, WH, MWK; Data Curation: LP, MB, LY; Writing: LP, LY, MWK; Supervision: LP, WH, LY, MWK.

## Acknowledgements

LP and MK were supported by NIH grant R01HD091846. We are grateful to Y.Galimov and O.Gusev for unpublished data on PIMT overexpression in *Daphnia* ephippia.

---

**Supplementary Fig. 1.**
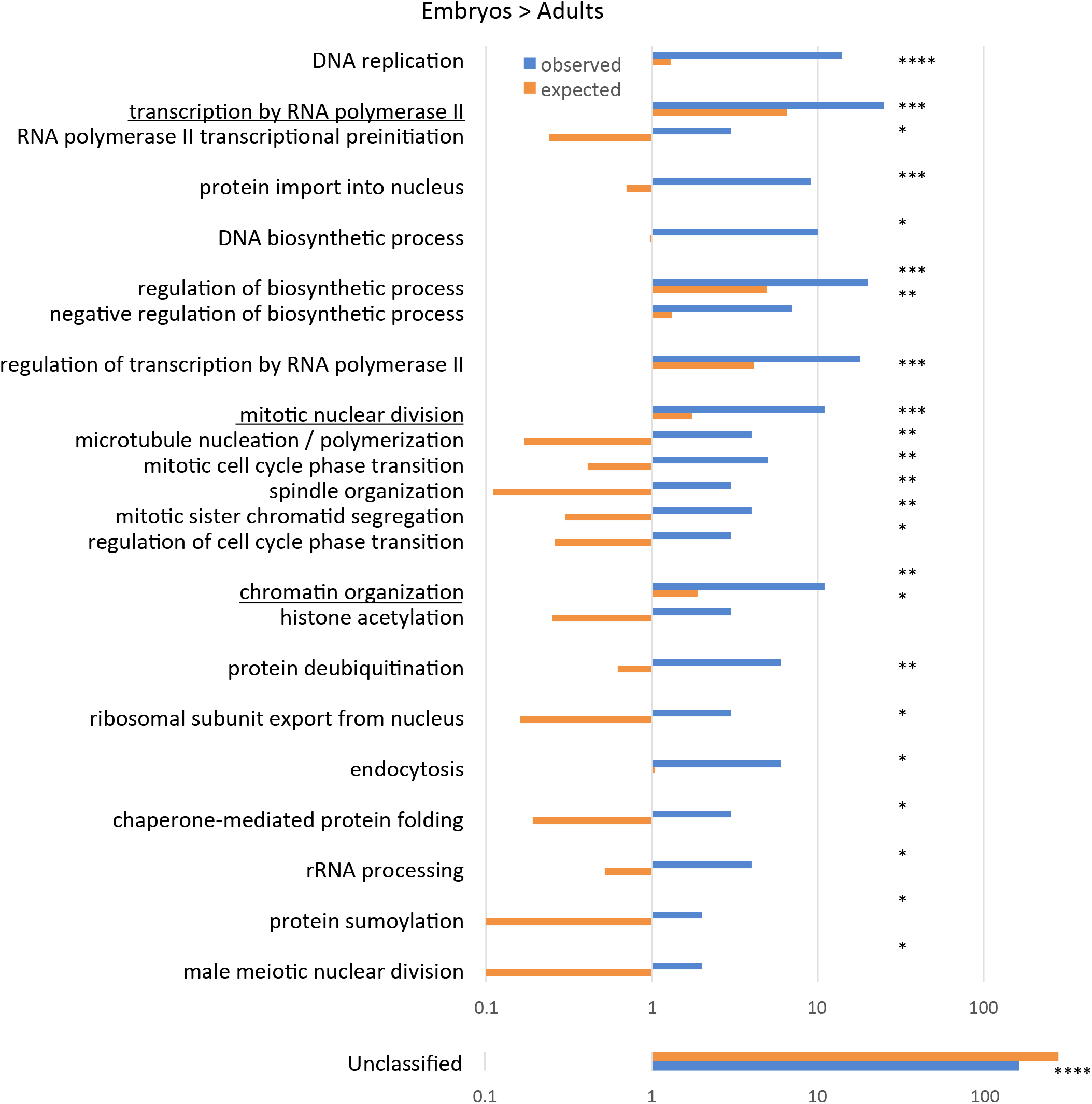
Overrepresentation of Biological Component gene ontologies with FDR<0.05 in the subset of proteins with higher expression (FDR<0.01) in subitaneous embryos vs. adults. Observed and expected number of proteins shown in the most specific GO term plus one parent GO up (underlined) if also significant. Overrepresentation FDR: **** <10^−6^; *** <10^−4^; ** <10^−2^; * < 0.05.

**Supplementary Fig. 2.**
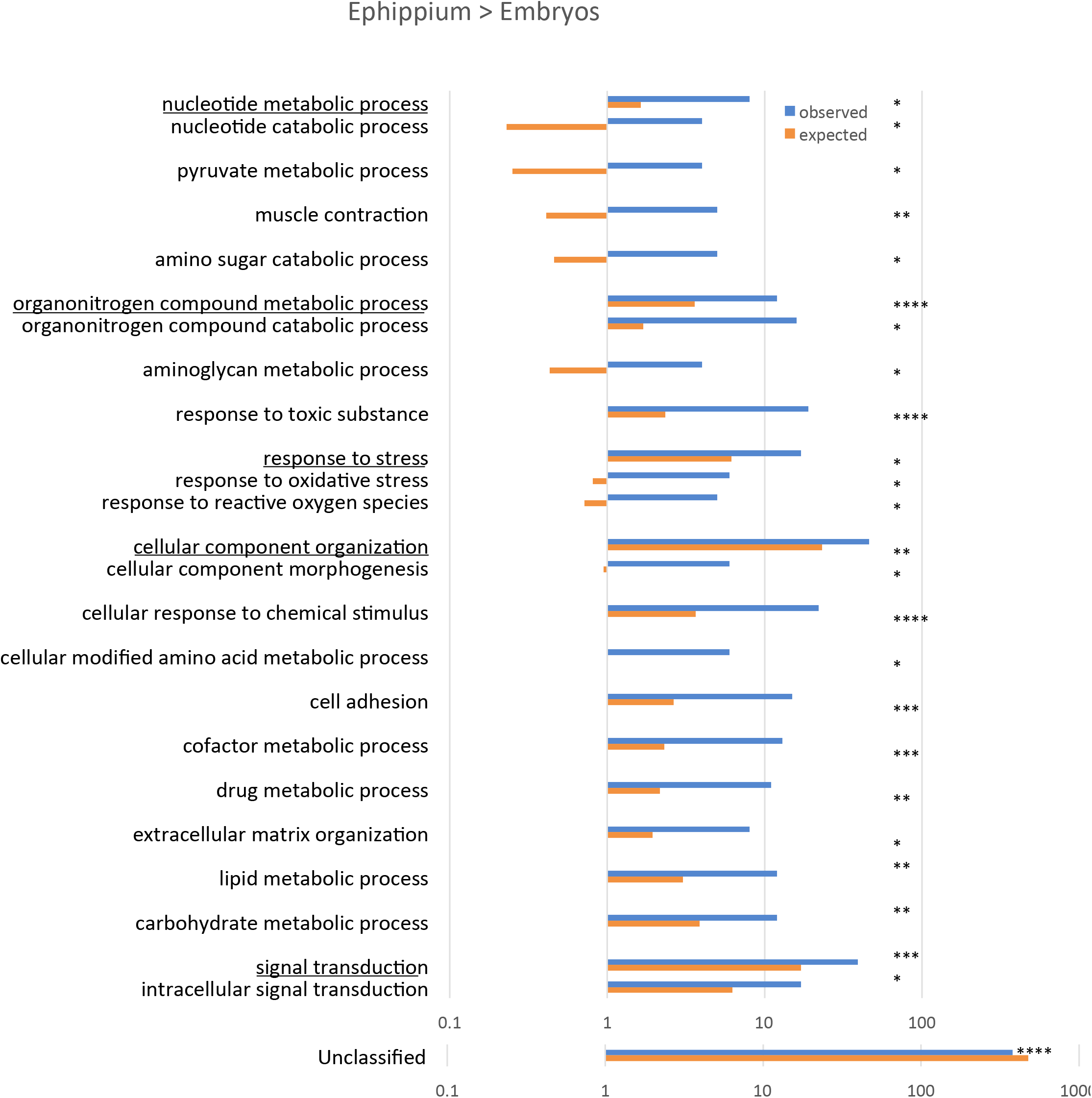
Overrepresentation of Biological Component gene ontologies with FDR<0.05 in the subset of proteins with higher expression (FDR<0.01) in the ephippium sample vs. subitaneous embryos samples. Observed and expected number of proteins shown in the most specific GO term plus one parent GO up (underlined) if also significant. Overrepresentation FDR as on Fig. S1.

